# The anti-phage mechanism of a widespread trypsin-MBL module

**DOI:** 10.1101/2025.09.03.673762

**Authors:** Pingping Huang, Jingxian Liu, Lijie Guo, Dongyang Xu, Lingbo Shen, Purui Yan, Chen Tong, Wenying Fei, Mengjun Cheng, Zhaoxing Li, Meiling Lu, Lei Zhang, Nannan Wu, Lianwen Qi, Yibei Xiao, Meirong Chen

## Abstract

Protease mediate activation of immune effectors is a conserved mechanism across life. This study identified a widespread and co-evolving trypsin-MBL module as a core effector in diverse bacterial immune systems (e.g., Hachiman, Argonaute, AVAST), protecting host via abortive infection. Focusing on Hachiman-trypsin-MBL, we demonstrated that, the protease activity of trypsin•HamAB is inhibited in the presence of ATP and MBL is an autoinhibited DNase with two inserted loops blocking the catalytic pocket. Upon infection, the HamB helicase senses invading nucleic acids and catalyzes the hydrolysis of ATP, activating the associated trypsin-like protease that cleaves specifically at two non-canonical KSS motif of MBL and releases its DNase activity. Cryo-EM structures of activated trypsin•HamAB bound with DNA reveal DNA binding and ATP hydrolysis lead to the release of the trypsin-like domain, enabling activation. Our work provides a comprehensive understanding on the mechanism of widely distributed trypsin-MBL module, wherein proteolytic activation of a toxic nuclease enables robust bacterial immunity while preventing self-toxicity through multi-layered control.

## Introduction

Across innate immune systems, the controlled activation of lethal effector molecules via proteolytic cascades represents a well-established mechanism^[1–5]^, exemplified by caspases in eukaryotic apoptosis^[5, 6]^. The discovery of analogous protease-based defense systems in bacteria—such as caspase-gasdermin homologs^[7, 8]^, SAVED-protease^[9–11]^, CRISPR-associated CHAT proteases^[12–15]^, TIR-Caspase systems^[16]^—has uncovered an ancient evolutionary strategy for the safe deployment of toxic immune effectors through proteolytic activation upon infection. As bacteria continue to evolve diverse defense mechanisms against invasive nucleic acids^[17–21]^, a central unresolved question concerns the full scope of immune processes governed by regulatory proteolysis and the diversity of effector substrates involved.

Previous studies have independently reported the frequent genetic co-occurrence of a trypsin-like serine protease gene adjacent to a gene encoding a metallo-β-lactamase (MBL) superfamily protein across multiple systems (e.g. ABC-3C, AVAST, and Retron)^[19,22–24]^, suggesting a potential functional partnership. However, the precise relationship within this trypsin-MBL module and its role in antiviral defense have remained unclear. Although bioinformatic analyses indicate that the MBL protein share structural homology with characterized enzymes such as metallo-β-lactamases, ComEC (a DNase), RNaseJ, and phosphorylcholine esterase (Pce), its biochemical activities and cellular functions still awaits to be discovered.

In this study, using systematic bioinformatics, we expanded the presence of trypsin-MBL modules to previously unrecognized systems, including those associated with Hachiman, Argonaute, Dsr2, and ABC-like components. We demonstrate that this module acts as a core effector in diverse defense systems, conferring anti-phage and immunity through abortive infection. Importantly, we show that MBL is a DNase auto-inhibited by two inserted loops that sterically block its catalytic pocket. The genetically linked trypsin-like protease serves as the initiator: upon detecting infection signals, it specifically cleaves these inhibitory loops, relieving suppression and unleashing DNase activity. In the Hachiman-associated trypsin-MBL system, this proteolytic activation is further secured by an additional layer of regulation wherein cellular ATP suppresses MBL cleavage. This inhibition is relieved through ATP hydrolysis by the helicase HamB upon sensing invading nucleic acids. Finally, we determined the Cryo-EM structure of the trypsin•HamAB in active state bound with 3′ overhang DNA, revealing that DNA-induced ATP hydrolysis enables the release and activation of the trypsin-like domain, which is further enhanced by higher-order oligomerization of trypsin•HamAB. Our findings elucidate the sophisticated defense mechanism of trypsin-MBL in which a protease activated nuclease effector cassette enables efficient immunity against invaders while avoiding toxicity through redundant autoinhibitory constraints, ensuring activation only upon genuine threat.

## Result

### The co-occurring trypsin-MBL is a widespread module in bacterial defense systems

Here, we performed a thorough bioinformatic analysis, and revealed over 1500 instances of trypsin–MBL pairs as candidate bacterial defense systems. Besides the reported association with defense systems AVAST, ABC-3C and Retron^[19, 22–24]^, in our analysis this trypsin-MBL module was also found frequently associated with the defense systems such as Hachiman^[18, 25, 26]^, short Argonaute^[27, 28]^, ABC-like^[22]^, and DSR2^[29]^ **(Fig.1A)**. Generally, the trypsin-like domain is fused at the N terminus of a component of the defense system, to which a gene annotated as MBL located adjacent **(Fig.1A)**. Phylogenetic analysis of MBL proteins showed clear clustering of trypsin fusions according to their respective systems **(Fig.1B)**, suggesting functional association and co-evolution between the trypsin and MBL components. This inference is further supported by the co-evolution analysis, which revealed a Spearman correlation of 0.677 between MBL and trypsin evolutionary distances **(Fig.1C)**. Sequence analysis predicted that the nuclease domain of HamAB and the PIWI domain of short Ago are predicted catalytically inactive. Furthermore, in other defense systems, the canonical effector domains are likely replaced by a trypsin-MBL module. We therefore propose that the trypsin-MBL may serve as the effector in these systems.

**Figure. 1.**
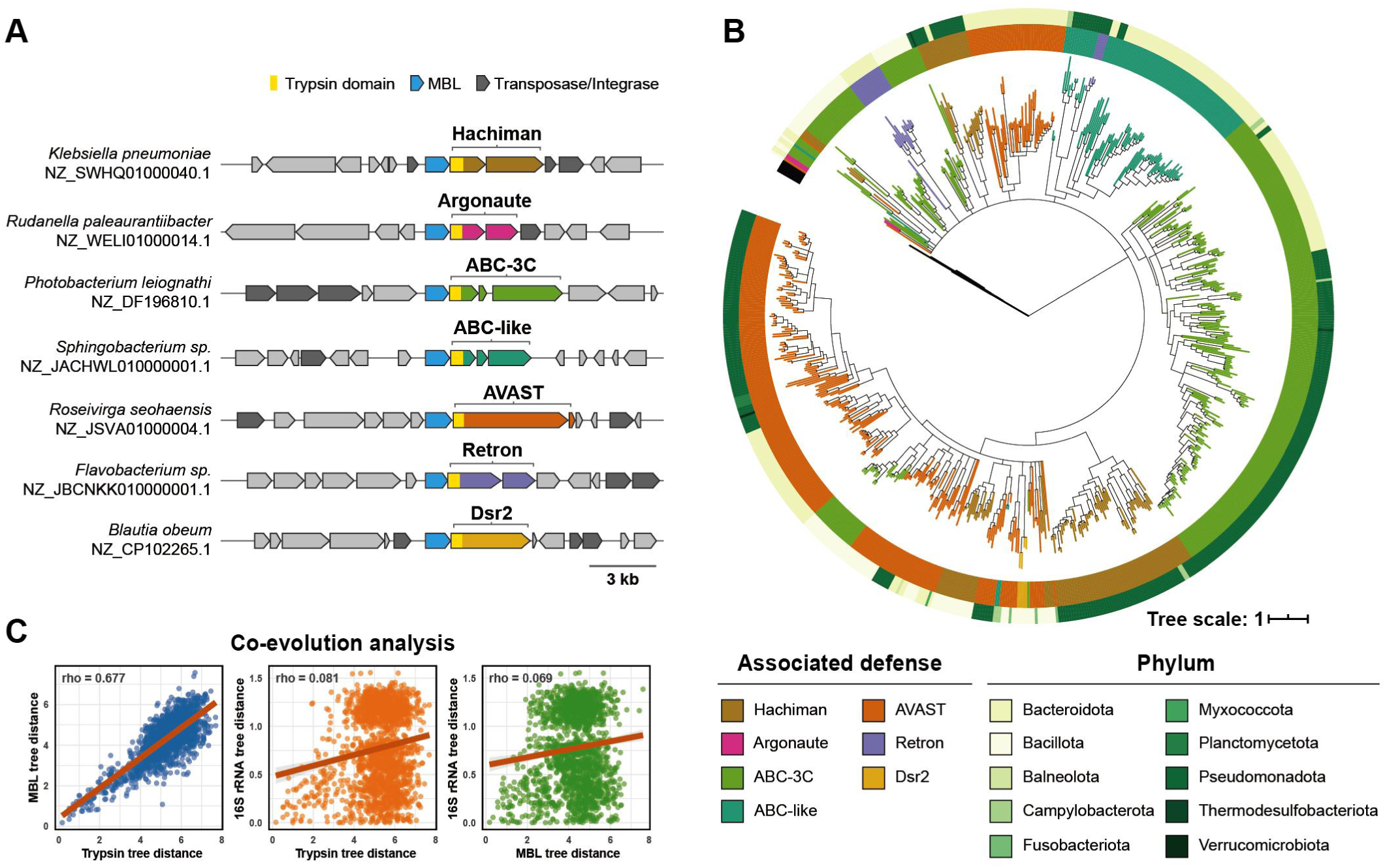
The co-evolving of a widespread trypsin-MBL module in bacterial defense systems. **(A)** Genomic contexts of representative defense-associated trypsin-MBL modules. The trypsin domains, MBLs, and transposase/integrases are colored yellow, blue, and grey, respectively. **(B)** Phylogenetic tree of MBLs associated with defense systems fused with trypsin-like domains. The inner circle indicates the associated defense system type; the outer circle indicates the corresponding bacterial phyla. **(C)** Co-evolution analysis of MBL proteins and associated trypsin domains with Spearman’s rank correlation. Pairwise distance matrix from the MBL phylogeny was compared to the corresponding matrix from the phylogeny of trypsin domains found in the same genomic locus. Comparisons with 16S rRNA phylogenetic distances served as negative controls. The graph depicts a random sample of 2000 data points for each correlation, with full dataset rho values reported.

### Trypsin-MBL confers bacterial anti-phage immunity

We next evaluated the bacterial defense function of several trypsin-MBL-containing systems, and successfully validated the bacterial defense function of AVAST-MBL, Hachiman-MBL, and Ago-MBL in *E.coli*. Expression of the AVAST-MBL system conferred resistance to 5 out of 12 tested *E. coli* phages **(Fig. 2A)**. Similarly, the Hachiman-MBL system exhibited strong anti-phage activity, protecting against 8 different phages. Both systems showed a strong preference for phages of the Myoviridae family **(Fig.2A)**.

**Figure 2.**
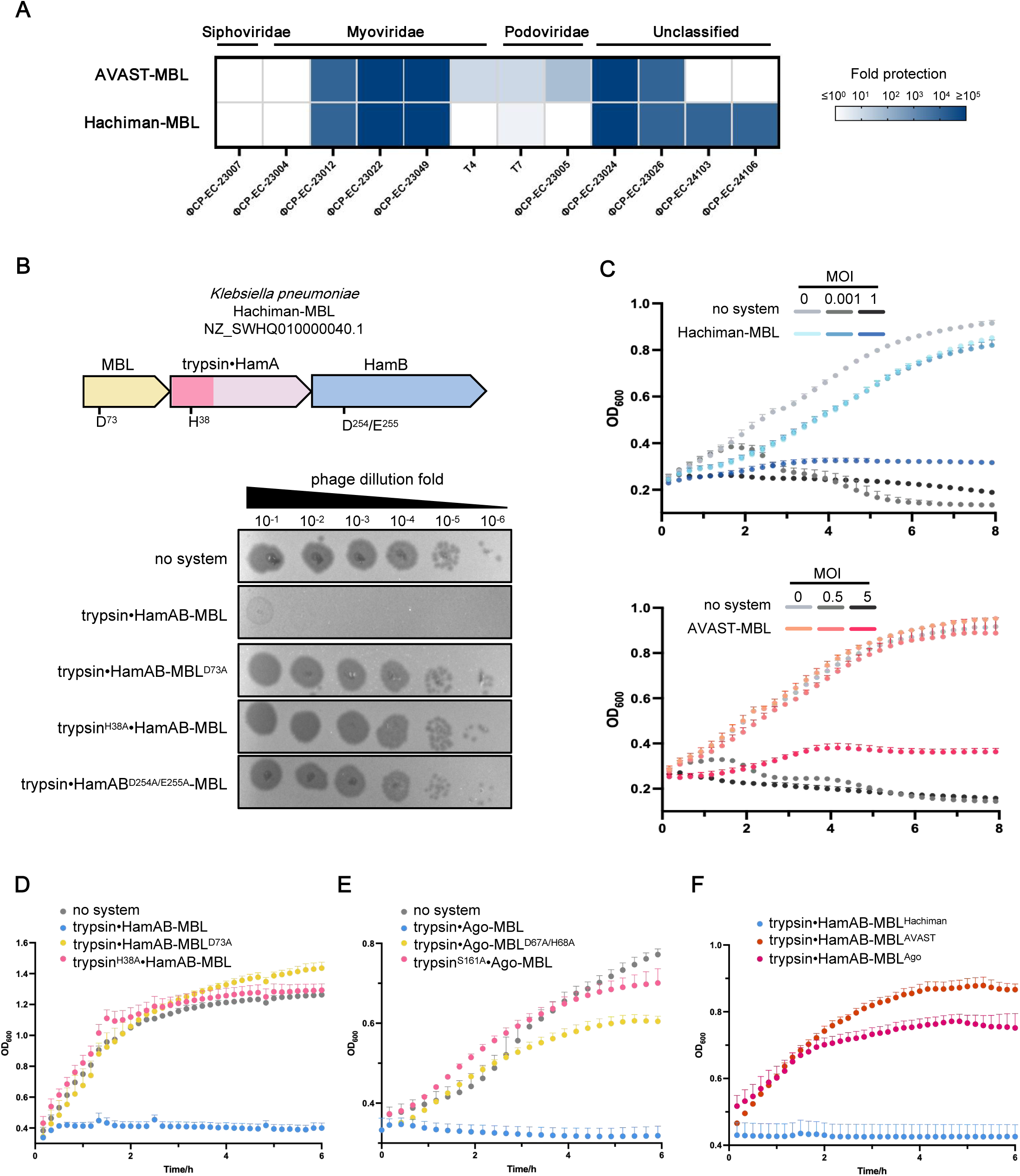
Trypsin-MBL confers anti-phage defense via abortive infection. **(A)** Phage protection mediated by *Vibrio cholera* AVAST–MBL and *Klebsiella pneumoniae* Hachiman–MBL systems expressed in *E. coli* MG1655, quantified as fold reduction in plaque formation via serial dilution assays. Phage titers were compared between strains harboring the defense system and a system-deficient control. **(B)** Plaque assay of phage ΦCP-EC-23022 on *E. coli* lawns expressing wild-type Hachiman–MBL or catalytic site mutants. A schematic of the operon with predicted catalytic residues is shown above. **(C)** Liquid infection assay of *E.coli* MG1655 expressing an empty vector or the Trypsin-MBL containing systems (Top: Hachiman-MBL, bottom: AVAST-MBL), after infection with phage ΦCP-EC-23022 at the indicated MOI. Error bars represent one standard deviation from three biological replicates. **(D, E)** Growth curves of *E.coli* expressing the trypsin-MBL systems (D. Hachiman-MBL, E. Ago-MBL) or their mutants, with a no-system strain as control. Curves show the mean ± SD of three replicates. **(F)** Growth curves of *E.coli* expressing the chimeric trypsin-MBL systems with MBL swapping between Hachiman, Ago, and AVAST.

Hachiman-MBL system comprises three components: a widely conserved HamB helicase, an HamA subunit that lacks the canonical nuclease catalytic residues but contains an N-terminal trypsin-like domain, and a MBL protein **(Fig.2B)**. The anti-phage defense of Hachiman-MBL required functional HamB, trypsin-like domain of HamA, and MBL, as mutation of predicted active sites in any of these proteins abolished anti-phage activity **(Fig.2B)**. Phage infection assays at different multiplicities of infection (MOI) revealed that cells expressing Hachiman–MBL exhibited limited growth arrest at low MOI but collapsed at high MOI—a hallmark of abortive infection **(Fig.2C)**, similar pattern was also observed with AVAST-MBL **(Fig.2C)**. Notably, even in the absence of phage, overexpression of HamAB and MBL induced by IPTG led to severe growth inhibition **(Fig.2D)**, and a similar toxic phenotype was also observed with Ago-MBL **(Fig.2E),** suggesting a potential anti-plasmid defense function. In both cases, toxicity depended on the catalytic residues of the trypsin and MBL domains **(Fig.2D,2E),** further suggesting that trypsin-MBL is a toxic effector module tightly linked to their catalytic activities. Furthermore, swapping MBLs between Hachiman, AVAST and Ago systems resulted in loss of toxicity **(Fig.2F)**, further supporting the hypothesis that trypsin-MBL function in a co-evolved and highly specific manner.

In conclusion, these findings establish the trypsin-MBL module as a core component of widespread bacterial defense systems. The defense function of these systems depends on the coordinated catalytic activities of both trypsin and MBL, and operates through an abortive infection mechanism.

### Hachiman-MBL is a nuclease-based bacterial defense system

To elucidate the role of the trypsin-MBL module in anti-phage defense, we investigated the biochemical function and mechanism of the Hachiman–MBL system from *Klebsiella pneumoniae*. To distinguish it from the canonical HamAB, the trypsin-containing form studied here was hereafter designated as trypsin•HamAB.

We firstly investigated the enzymatic activities of trypsin•HamAB proteins. The ATPase activity of the trypsin•HamAB helicase was assessed using various nucleic acid substrates including ssDNA, ssRNA, dsDNA, plasmid DNA, duplex DNA with either 3′ or 5′ overhang. The results demonstrated that all forms of DNA tested—but not ssRNA—stimulated ATP hydrolysis, with duplex DNA with 3′ overhangs eliciting the strongest activation **(Fig.S1A)**. Subsequent DNA unwinding assays using partially duplex DNA substrates containing either 5′ or 3′ single-stranded overhangs revealed that trypsin•HamAB exhibits robust ATP-dependent helicase activity specifically toward 3′ overhang DNA, while with minimal activity on 5′ overhangs **(Fig.S1B)**. Together, the ATPase and helicase activities indicate that DNA bearing a 3′ overhang serves as the preferred substrate for trypsin•HamAB. We also evaluated the nuclease activity of trypsin•HamAB against various nucleic acids substrates—including plasmid DNA, ssDNA, dsDNA, and ssRNA—and found no detectable nuclease activity even in the presence of 3′ overhang DNA **(Fig.S1C)**.

The MBL protein found in Hachiman and other trypsin-MBL containing systems exhibits structural homology to metal-β-lactamases, ComEC (which possesses DNase activity), RNaseJ and phosphorylcholine esterase (Pce)^[30–34]^. Therefore, we test these activities of MBL using a panel of substrates including β-lactam, plasmid DNA, ssDNA, dsDNA, RNA, or choline. While MBL alone is inactive to any of the above substrates **(Fig.3A, S2A-S2C)**, surprisingly, the incubation of MBL with trypsin•HamAB resulted in robust DNase activity against plasmid **(Fig.3A)**, ssDNA, and dsDNA **(Fig.S2A)**, but not toward β-lactam, RNA, or choline **(Fig.S2A-S2C)**. Notably, this DNase activity was dependent on both MBL and the trypsin-like protease, as mutations in the active site of either component abolished DNA cleavage activity **(Fig.3A)**.

**Figure 3.**
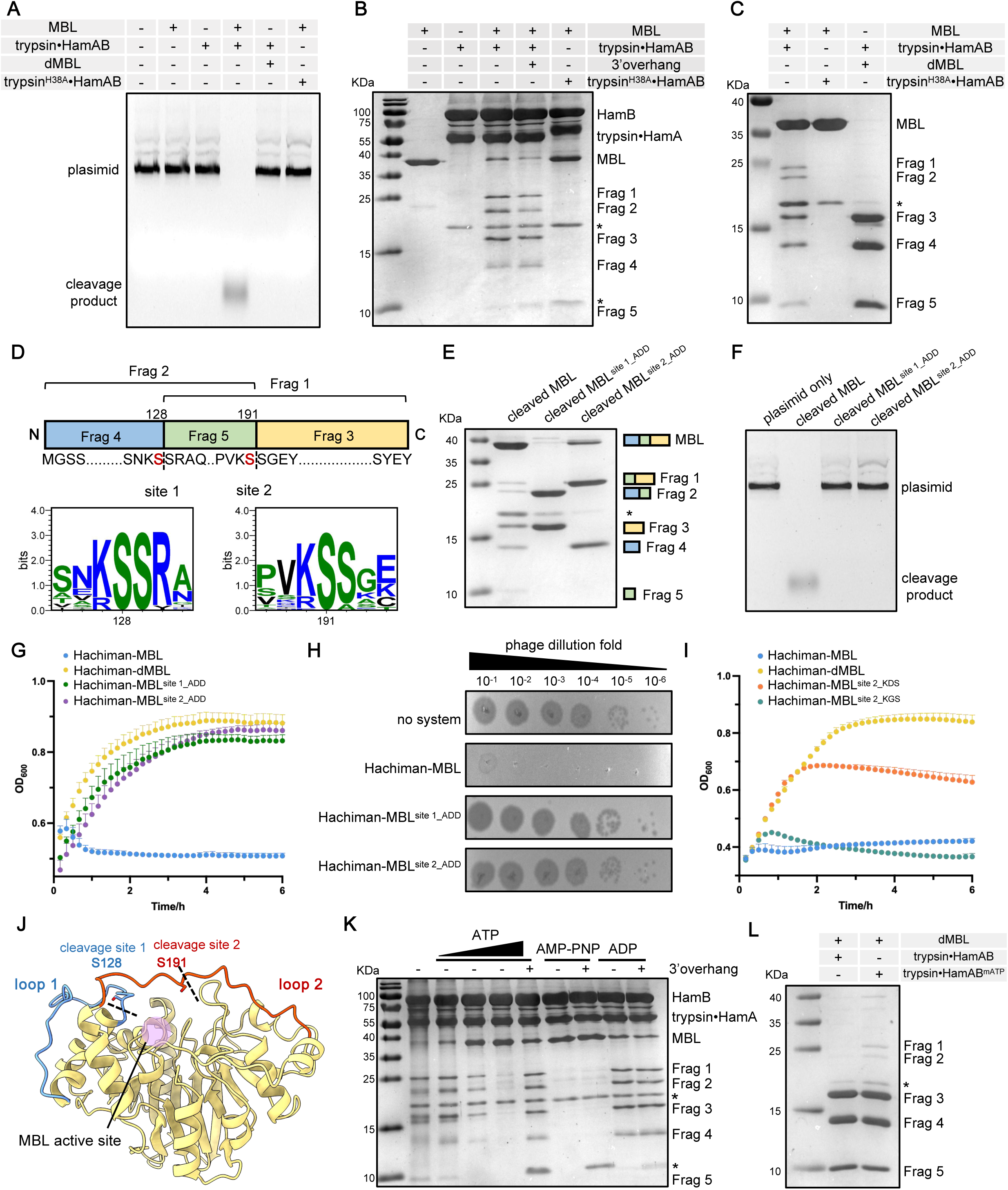
trypsin•HamAB activates DNase activity of MBL by specifically cleaving the autoinhibitory loops of MBL. **(A)** Plasmid cleavage assay of trypsin•HamAB-MBL analyzed by urea-PAGE. **(B)** Cleavage of MBL by trypsin•HamAB *in vitro*. The cleavage products are labeled as Frag 1-5. A band of impurities is indicated with an asterisk. **(C)** *In vivo* cleavage of MBL by trypsin•HamAB. The trypsin mutant HamA^H38A^ was used as negative control. The cleaved MBL was pulled down from cultures co-expressing MBL and trypsin•HamAB and analyzed by SDS-PAGE. dMBL: catalytic inactive MBL mutant. **(D)** Mapping of MBL cleavage sites by mass spectrometry. The three fragments (Frag 3-5) from (C, lane 3) were sequenced. Cleavage sites are mapped onto the primary structure of MBL(top). N- and C-terminal sequences of each fragment are aligned below, with cleavage site serines highlighted in red. The Fragment 1-5 corresponding to the SDS-PAGE of (B,C) was indicated. The conservation analysis of the cleavage site among short-form MBL is shown at the bottom. **(E)** Validation of site-specific cleavages using MBL mutants with KSS motif substituted to ADD residues. Cleavage fragments are schematized beside the gel. **(F)** Validation of site-specific cleavage induced activation of MBL by plasmid cleavage assay. **(G)** Growth curves of *E.coli* expressing wild-type Hachiman-MBL or the cleavage sites mutants. **(H)** Plaque assay of phage ΦCP-EC-23022 on *E. coli* expressing wild-type Hachiman–MBL or cleavage site mutants. **(I)** Cleavage site preference of trypsin•HamAB analyzed by growth toxicity. **(J)** The cleavage sites mapping onto the inhibitory loops (blue and red) of MBL structure predicted by AF3. The active site is labeled with pink sphere. **(K)** Inhibition of MBL cleavage by ATP and its analogs, in the presence or absence of 3’overhang DNA. **(L)** *In vivo* cleavage of MBL by trypsin•HamAB bearing ATP-binding site mutations. (trypsin•HamAB^mATP^: trypsin•HamAB with mutations G158A/K159A/S160A/D254A/E255A).

Together, these findings indicate that Hachiman-MBL functions as a DNA-specific nuclease system in which trypsin•HamAB activates the nuclease activity of MBL.

### Trypsin•HamAB activates MBL by specifically double cleaving the inhibitory loops

Structure alignment of the AF3-predicted MBL model with its homologs revealed a significant structural divergence: MBL contains a long inserted loop (184–203 aa) that extends directly over the catalytic pocket **(Fig.S2D)**. This inserted loop is absent in metallo-β-lactamases, ComEC, Pce, and RNaseJ^[32–34]^ **(Fig.S2E)**, but is structurally conserved across trypsin-MBL-containing systems **(Fig.S2D)**. We hypothesize that this loop may sterically occlude substrate access to the MBL active site, thereby render MBL inactive. Given the conserved genetic linkage between MBL and the trypsin-like domain within HamAB, together with the finding on trypsin-dependent activation of MBL **(Fig.3A)**, we speculate that, upon phage infection, the trypsin-like domain may cleave the inserted loop, relieving autoinhibition and activating the DNA-cleaving function of MBL. Notably, this trypsin-induced activation of MBL may represent a general mechanism employed by trypsin-MBL-associated systems.

To elucidate the mechanism how the ATP-dependent helicase trypsin•HamAB activates MBL, we incubated MBL and trypsin•HamAB in buffer containing 2 mM ATP, with or without 3′ overhang DNA, and analyzed the samples by SDS-PAGE. Incubation with trypsin•HamAB resulted in a pronounced reduction of full-length MBL and the appearance of five lower-molecular-weight bands (Frag 1-5) **(Fig.3B)**. This proteolysis was dependent on trypsin-like activity of trypsin•HamA and was enhanced in the presence of 3′ overhang DNA **(Fig.3B)**, suggesting that both trypsin•HamA and HamB are required for efficient cleavage of MBL. These findings support a model in which DNA unwinding by HamB induces activation of trypsin-like domain within HamA, enabling proteolytic processing of MBL.

We further examined MBL processing under *in vivo* conditions. Pull-down of MBL from cells co-expressing trypsin•HamAB and MBL revealed a similar cleavage pattern **(Fig.3C)**. Remarkably, the catalytically inactive mutant MBL^D73A^ (dMBL), was completely processed compared to tthe wild-type MBL, yielding three specific fragments (Frag 3-5) **(Fig.3C)**. This enhanced processing likely occurred due to the absence of toxicity that allowed uninterrupted hydrolysis.

Notably, the purified cleaved MBL alone exhibited strong plasmid-degrading activity **(Fig.S2F)**, confirming that proteolytic processing releases its DNase function, which is responsible for both cellular toxicity and anti-phage defense. This result also indicates that the cleaved active MBL alone is able to digest plasmid without the involvement of the helicase activity of trypsin•HamAB. These findings further support a model in which trypsin•HamAB acts as a sensor and activator, while MBL cleaved by trypsin•HamAB serves as the effector DNase.

To identify the exact cleavage sites on MBL essential for releasing its DNase activity, the three proteolytic fragments were analyzed by mass spectrometry sequencing. All three fragments were mapped to MBL and collectively spanned the full-length protein **(Fig.3D)**. Cleavage occurred specifically between two serine residues at S128–S129 and S191–S192 **(Fig.3D)**, diverging from the canonical lysine/arginine recognition pattern of trypsin^[35]^. The non-canonical cleavage specificity of trypsin-like domain was further verified using a synthetic fluorescence-labeled peptide containing a KR motif, which remained uncleaved in contrast to the positive control treated with commercial trypsin **(Fig.S1D)**. Interestingly, a lysine residue was present at the P2 position in both cleavage sites **(Fig.3D)**, forming a conserved KSS motif.

The specificity of proteolytic processing was further confirmed through mutational analysis. Introduction of ADD mutations at either the ^127^KSS^129^ or ^190^KSS^192^ site rendered MBL resistance to cleavage at corresponding site, resulting in the disappearance of fragment 1 or 2 (intermediate products) and fragment 5 (the central fragment between the two cleavage sites) **(Fig.3E)**. Interestingly, disruption of single cleavage site will be sufficient to abrogate MBL’s DNase activity **(Fig.3F)**, and thus eliminate the toxicity of MBL effector **(Fig.3G)** and its Abi-based anti-phage activity **(Fig.3H)**. These results confirmed that dual-cleavage at two KSS motifs is required for activation of Hachiman-MBL defense system.

The residue preference at P1 site by trypsin-like is further investigated. We mutate serine into bulky aspartic acid and small glycine, respectively, and examine their effect on MBL cleavage by cell toxicity assay. The results showed that while the larger aspartic acid substitution would significantly relieve cell toxicity of MBL, the glycine of similar size as serine did not affect the cleavage of MBL, thus remained cell growth arrest as wild type MBL **(Fig.3I)**. These findings underscore that trypsin-mediated cleavage of MBL prefers small residues at P1 site, such as serine and glycine.

Mapping the cleavage sites onto the predicted structure of MBL elucidated the mechanism by which dual proteolytic cleavage releases its DNase activity. The cleavage site at S191 is just situated within the previously described auto-inhibitory loop (184–203 aa) that directly occludes the catalytic pocket, while the second site at S128 lies on an adjacent loop (107–129 aa) **(Fig.3J)**. Cleavage at both sites thus appears to release steric hindrance over the catalytic pocket, enabling large DNA substrate access. Both loops are absent in other MBL homologs—including the functional DNase ComEC—but are conserved in trypsin-MBL-containing systems **(Fig.S2D, S2E)**, suggesting a specific auto-inhibitory role tailored for trypsin-dependent activation of MBL.

Notably, phylogenetic analysis and sequence alignment categorize MBL proteins within Hachiman-MBL systems into two distinct groups **(Fig.S3A, S3B)**: one group, as studied here, retains highly conserved KSS motifs in both loops **(Fig.3D, Fig.S3B)**, while the other group featuring longer auto-inhibitory loops lacks the KSS signature but contains a conserved IS(G)N motif **(Fig.S3B)**. This IS(G)N motif, recently identified on bioRxiv, has been confirmed as the cleavage sites in long-form MBL^[36]^. Although trypsin-mediated proteolytic processing occurs in both groups, the P1 residue preference differs—serine in the first group and isoleucine in the second. To test functional compatibility, we performed MBL swaps between the two groups. While MBL exchange within the same group (KSS type) remains functional that causes cell growth arrest, exchange between the two groups impedes the defense function as it significantly alleviates cellular toxicity **(Fig.S3C)**, further supporting co-evolution between trypsin and its cognate MBL substrate.

Collectively, these findings demonstrate that the two insertion loops inhibit MBL activity, and that trypsin-mediated cleavage at specific sites within these loops relieve this repression, thereby activating DNase function.

### ATP inhibits cleavage and activation of MBL

We found that in the absence of exogenous triggers, trypsin•HamAB alone can cleave MBL **(Fig.3B, Lane 3)**, suggesting the existence of a potential auto-inhibition mechanism within the cell. While analyzing the proteolysis of MBL, we attempted to enhance hydrolysis by supplementing additional DNA and ATP, both essential components of the antiviral signaling pathway mediated by HamAB. Unexpectedly, MBL cleavage by trypsin•HamAB was increasingly inhibited in the presence of 0.5 to 5 mM ATP **(Fig.3K)**. This observation led us to hypothesize that ATP might suppress the innate proteolytic activity of the trypsin-like domain by binding to the ATPase pocket of HamB. We therefore assessed MBL cleavage by trypsin•HamAB in the absence of ATP and found that trypsin•HamAB alone hydrolyzes MBL much more efficiently **(Fig.3K)**, supporting ATP-mediated inactivation of the trypsin•HamAB–MBL. Notably, this inhibition could be reversed by the addition of trigger 3′ overhang DNA.

To investigate whether the hydrolysis of inhibitory ATP is required for releasing inhibition, we tested the non-hydrolyzable analog AMP-PNP. Similar to ATP, AMP-PNP inhibited MBL cleavage; however, this inhibition could not be relieved by 3′ overhang DNA, indicating that relief of inhibition depends on DNA-triggered ATP hydrolysis**(Fig.3K)**. Consistent with this, ADP did not inhibit cleavage **(Fig.3K)**.

The inhibitory role of ATP aligns with our earlier findings and helps clarify the regulatory scenario: in the absence of infection, the physiological concentration of cellular ATP would inhibit trypsin•HamAB, which can be reversed specifically by invading DNA that induces ATP hydrolysis. Interestingly, the ATP binding and hydrolysis site of HamB is dispensable for MBL cleavage *in vivo* **(Fig.3L)**, implying that ATPase activity may be designated for removing ATP inhibition. Notably, in a very recent study the ATPase-independent effector activity has also been observed in many helicase-nuclease mediated defense systems—including Gabija, canonical Hachiman, AbpAB, and Mokosh II—in which loss of ATPase activity does not compromise effector function *in vitro* with unknown mechanisms^[37, 38]^. Our study may offer mechanistic insights into the role of ATPase in these defense systems, as it may regulate the inhibition and activation of the effector. These consistencies suggest that an ATP-mediated auto-inhibitory strategy is likely widespread among ATPase-associated bacterial defense systems.

Taken together, our study reveals that cellular ATP would inhibit trypsin•HamAB to avoid toxicity, which can be released via ATP hydrolysis when sensing invading DNA.

### The structures of active trypsin**•**HamAB bound with DNA reveals the release of trypsin for activation

To elucidate the mechanism by which DNA activates trypsin-like activity in trypsin•HamAB for subsequent MBL cleavage, we determined the cryo-EM structures of the active form of trypsin•HamAB by incubating it with 3′-overhang DNA and ATP. These structures reveal multiple oligomeric assemblies, including a heterodimeric trypsin•HamAB-DNA complex, a dimer of heterodimers bound to DNA, and a trimer of heterodimers bound to DNA, at a resolution of 3.2, 3.2, 3.5 Å, respectively, with a particle ratio of approximate 1:3:2 **(Fig.4A, 4B, Fig.S4A, S4B)**.

**Figure 4.**
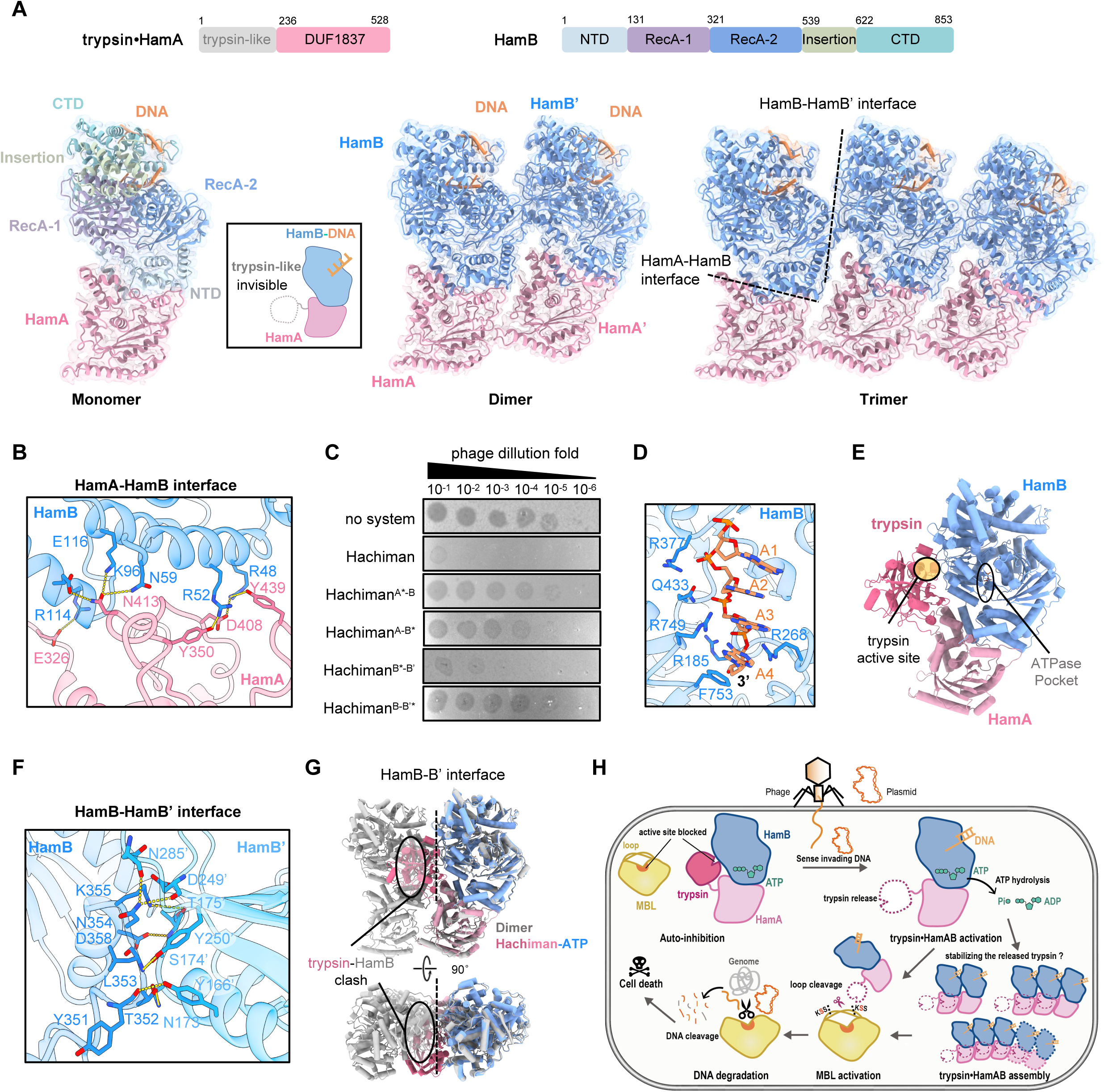
The structures of activated trypsin•HamAB. **(A)** Cryo-EM structures of trypsin•HamAB shown in cartoon representation. Domain coloring corresponds to the schematic diagram shown on top. Structures include a DNA-bound trypsin•HamAB, heterodimer bound with DNA (left), dimer of heterodimer bound with DNA (middle), trimer of heterodimer bound with DNA (right). The trypsin-like domain is disordered in all the structures due to flexibility and indicated with a dashed circle in the inset. **(B)** The interactions at HamA-HamB heterodimer interface. **(C)** Plaque assay of phage ΦCP-EC-23022 on *E.coli* lawns expressing wild type Hachiman-MBL or the oligomerization interface mutants. **(D)** The nucleotides of 3’overhang DNA embraced in HamB helicase channel.**(E)** AF3-predicted model of trypsin•HamAB in complex with ATP. The active sites of ATPase and trypsin are indicated. **(F)** Interactions at the B-B’ interface in higher-order oligomers of trypsin•HamAB. **(G)** Superimposition of inhibited trypsin•HamAB-ATP onto active trypsin•HamAB-DNA. **(H)** The proposed mechanism of Hachiman-MBL defense system.

In trypsin•HamA, the N-terminal trypsin-like domain is invisible, while the C-terminal domain (236–528 aa), named DUF1837, consists of a central four-stranded β-sheet flanked by α-helices on both sides **(Fig.4A)**, reminiscent of the core structural motif in the PD-(D/E)XK phosphodiesterase superfamily^[39]^. Notably, the catalytic residues are missing in trypsin•HamA active site, consistent with the lack of nuclease activity **(Fig.S1C)**. HamB, classified within the SF2 helicase family^[40]^, possesses the characteristic RecA1 and RecA2 domains with an insertion, along with N-terminal and C-terminal domains exhibit a predominantly α-helical structure **(Fig.4A)**.

Within the trypsin•HamAB heterodimer, the C-terminal domain of HamA engages in extensive electrostatic interactions with the N-terminal α-helical structure of HamB burying a surface area of 1694 Å². At the primary interface, R48 and R52 of HamB form hydrogen bonds with Y350, D408, and Y439 of HamA, while the side chains of N59 in HamB hydrogen-bond with the main-chain atom of N413 of HamA. Additionally, K96, R114, and E116 in HamB participate in electrostatic interactions with the side chains of N413 and E326 in HamA **(Fig.4B)**. Mutations of key AB interfacial residues Y350A/D408A/Y439A or N59D would significantly abrogated phage protection **(Fig.4C: Hachiman^A*-B^, Hachiman^A-B*^)**. Density corresponding to DNA was observed within the helicase channel of HamB. The phosphate backbone of four nucleotides from the 3′ overhang could be traced, with the first nucleotide confidently modeled. The backbone mediates extensive electrostatic interactions with HamB, engaging positively charged residues including R185, R268, R377, Q433, and R749. Additionally, the first nucleotide stacks with F753 **(Fig.4D)**. As expected, no ATP density was detected in the ATPase pocket of HamB, consistent with an activated state in which DNA binding stimulates ATP hydrolysis.

The fully disordered trypsin-like domain in this activated trypsin•HamAB indicates its high flexibility after activation **(Fig.4A)**. We therefore speculate that the DNA binding—induced ATP hydrolysis—triggers the release and activation of trypsin-like domain. Supporting this, an AF3-predicted model of trypsin•HamAB in complex with ATP showed the trypsin-like domain restrained at the HamAB heterodimer interface—closely access to the ATPase pocket **(Fig.4E)**. Interestingly, the trypsin orientates with the catalytic pocket facing to the interface, representing a substrate-inaccessible state. The comparison of active state and inhibited state of trypsin•HamAB reveals the conformational change of trypsin from sequestered to flexible. Therefore, these observations raise an idea that ATP hydrolysis at the interface of HamAB may disrupt their association with trypsin-like domain, and thus release and activate it. Notably, the inhibition of fused effector due to the association with HamAB has also been observed in previously reported Cap4-HamAB structure **(Fig.S5)**^[26]^, further supporting our trypsin•HamAB inhibition and activation model.

Monomeric trypsin•HamAB-DNA units further oligomerize through angled stacking, forming a fan-like architecture**(Fig.4A)**. Particles containing up to five units were observed during 2D classification, despite of weaker peripheral densities due to alignment flexibility **(Fig.S4A)**. The oligomerization interfaces are flat and extended, primarily mediated by HamA-HamA′, HamB-HamA’ and HamB-HamB′ contacts **(Fig.4F)**. The HamB-HamB′ interface mainly contributes to oligomerization through electrostatic interactions involving residue Y166, E169, N173, S174, T175, D249, Y250, and N285 from one HamB protomer with residue Y351, T352, L353, N354, K355, D358 from another **(Fig.4F)**. Same as in trypsin•HamAB heterodimer, the trypsin-like domain of all the HamA protomers in dimer or trimer of heterodimer are invisible. Strikingly, superimposition of predicted inhibited trypsin•HamAB onto the trypsin•HamAB in oligomeric state reveals steric clashes between the trypsin-like domain and a HamB protomer from neighboring unit **(Fig.4G)**. We speculate that the oligomerization via HamB-HamB′ interface may occupy the binding site used by the inhibited trypsin-like domain, thereby stabilizing the released active conformation. Consistent with this observation, introducing multiple mutations (Y166A/N173A/S174W/T175W/Y250A or T352A/N354A/K355W/D358W) at the HamB-HamB′ interface yielded functional proteins but impaired bacterial defense against phage infection **(Fig.4D: Hachiman^B*-B’^, Hachiman^B-B’*^)**, underscoring the importance of trypsin•HamAB oligomerization in Hachiman-MBL mediated anti-phage defense.

In summary, we proposed that DNA binding to trypsin•HamAB induces ATP hydrolysis, leading to the release and activation of the sequestered trypsin-like domain for MBL cleavage, which is likely enhanced by oligomerization of trypsin•HamAB **(Fig.4H)**.

## Discussion

In this study, we elucidate the function and mechanism of a co-evolving trypsin-MBL module involved in bacterial defense. This system represents a unique form of protease-mediated activation of a nuclease through site-specific proteolysis, conferring anti-phage capability to the host via an abortive infection mechanism.

Unlike its intrinsically active homologs, MBL is an auto-inhibited nuclease that employs loop insertions to block its catalytic pocket. This structural arrangement serves as a regulatory mechanism, enabling strict control of the toxicity by the upstream trypsin-like domain. The trypsin-like domain recognizes a KSS motif within the inhibitory loop and specifically cleaves after the first serine—rather than after lysine, which is the canonical residue targeted by conventional trypsin. Although this cleavage pattern has not been previously documented, the preference for serine is reminiscent of elastase, which favors residues with small side chains (e.g., Ala, Gly, Ser)^[41, 42]^. The trypsin-like domain predicted by AF3 overall adopts a fold similar to that of canonical trypsin. However, detailed examination of its catalytic pocket reveals notable differences: whereas canonical trypsin typically contains two glycine residues at the entrance of the S1 pocket—accommodating long side chains such as those of arginine or lysine^[43]^—the trypsin-like domain studied here exhibits substitutions with larger residues (Thr and Cys), analogous to the Thr and Val residues found in elastase^[44]^. These substitutions result in a shallow and narrow S1 pocket that likely restricts access to residues with small side chains, probably explaining the preference cleavage after serine. Consistently, mutation of serine to the bulky aspartic acid, but not to the small glycine, attenuated the toxicity mediated by cleaved MBL. Together, these findings indicate that the trypsin-like domain in HamA both structurally and functionally resembles elastase rather than canonical trypsin. Notably, the specific recognition and cleavage of MBL by its cognate trypsin is further supported by both the evident co-evolution of the trypsin-MBL pair and the functional impairment observed upon swapping MBL across distinct systems (e.g., Hachiman, Ago, or AVAST), which results in loss of antiviral defense.

Interestingly, the activation of MBL necessitates dual cleavages at two spatially distant loops, whereas other known protease-associated defense systems—such as those mediated by CHAT or CalpS protease^[9–15]^—typically activate their substrates via a single proteolytic event. Although this requirement for two precise incisions may appear inefficient, both anti-phage assays and toxicity assessments demonstrate that the trypsin-MBL system exhibits robust defensive capability. We propose that the dependence on site-specific dual cleavages for MBL activation may represent a protective strategy to minimize the risk of non-specific digestion by endogenous cellular proteases, thereby preventing unintended activation and conferring avoidance of auto-toxicity in the absence of phage infection.

Unlike the Hachiman system, which degrades DNA through a single-step activation process where invading DNA directly triggers the HamA nuclease, the Hachiman-MBL system employs a two-step mechanism involving sequential activation of a trypsin-like domain of HamA and the MBL nuclease. This multi-level activation offers significant advantages. Firstly, the enzymatic cascade enables substantial signal amplification, facilitating rapid dissemination of the defense signal throughout the cell—analogous to the cascade initiated by initiator and effector caspases during apoptosis^[5]^. Secondly, the indirect nature of this activation allows for more precise tuning of immune responses, particularly through auto-inhibitory regulation, which helps minimize potential toxicity to the host cell and ensures targeted activation only upon specific infection cues.

In addition to the auto-inhibitory insertion loops within MBL, our findings further identify cellular ATP as a critical metabolite that exerts a second layer of allosteric regulation by suppressing the activation of trypsin•HamAB. This ATP-mediated inhibition represents a sophisticated regulatory strategy: as one of the most abundant cellular molecules, ATP is drastically consumed for viral replication during phage infection, serving as an ideal regulator fine-tune the inhibition-activation of the defense systems. Indeed, ATP-mediated inhibition has also been previously observed with another ATPase-associated system

Gabija and Her-Sir2^[37, 45]^. These together support the idea that ATP-mediated inhibition probably represents a universal mechanism in antiviral immunity Synergistically, other ATP-depleting defense systems, such as RADAR^[46, 47]^ [46–47], CRISPR-CAAD^[48, 49]^ and PNP-containing systems^[50]^, may also facilitate Hachiman-MBL activation, thereby promotes the antiviral immunity.

## Acknowledgments

We would like to thank the Instrument Analysis Center (IAC) at Shanghai Jiao Tong University for Cryo-EM data collection.

## Author contributions

CMR and XYB conceived the project.

CMR, XYB, QLW, WNN supervised the project.

LJX, HPP, SLB, YPR performed protein purification, biochemical assays, and cell-based assay.

HPP, LJX, LZX determined the Cryo-EM structure. GLJ performed the phage assays.

XDY performed bioinformatics analyses.

CMR, XYB, QLW, WNN, HPP, GLJ, LJX, SLB, TC, FWY, CMJ, LML, ZL analyzed data.

CMR wrote the original draft.

CMR, XYB and XDY reviewed and edited the manuscript. All authors discussed the results and contributed to the final manuscript.

## Data availability

Atomic coordinates, maps and structure factors of the reported cryo-EM structures have been deposited in the Protein Data Bank under accession numbers 9WH1 (DNA-bound HamAB heterodimer), 9WHK (DNA-bound dimer of HamAB heterodimer), 9WHU (DNA-bound trimer of HamAB heterodimer). and in the Electron Microscopy Data Bank under accession codes 65964, EMD-65968, and EMD-65977

## Methods

### Phylogenetic analysis

The genomic sequences and genome annotations of bacteria annotated genomes were retrieved from the RefSeq database^[1]^. Protein sequences were searched against the PF13365 HMM profile using hmmscan from HMMER (v3.3.2)^[2]^ with default parameters to identify Trypsin_2 domain containing proteins. For each of the Trypsin_2-containing proteins, the immediate upstream and downstream neighboring proteins were extracted and screened for MBL homology using the BLASTP program from the NCBI-BLAST+ package (v2.16.0)^[3]^ with an E-value cutoff of 0.001. The identified trypsin-MBL pairs were subsequently examined for defense system association using DefenseFinder (v2.0.1)^[4]^. The phylogenetic analysis was conducted for the MBL proteins in the defense-associated trypsin-MBL pairs. Protein sequences were clustered using MMseqs2 easy-cluster (release 17-b804f; --min-seq-id 0.9 -c 0.9 --cov-mode 1)^[5]^ to remove redundancy and define representative sequences. Multiple sequence alignments were generated using MAFFT (v7.526; -ep 0 --genafpair --maxiterate 1000)^[6]^, with RNase Z (n=5) as the outgroup. Alignments were trimmed with trimAl (v1.5.rev0; -gt 0.2 -cons 70)^[7]^ to remove gaps and poorly conserved sites. Phylogenetic tree was constructed using the IQ-TREE program (v3.0.1; -m MFP -nm 500 -bb 1000)^[8]^, with model selection based on Bayesian Information Criterion (Best-fit model: Q.PFAM+F+I+R10). Phylogenetic tree was visualized with iTOL (v7; itol.embl.de)^[9]^, where branches with <70% bootstrap support were collapsed into polytomies.

### Co-evolution analysis

To investigate the co-evolutionary relationship between the trypsin domains and MBL proteins, we selected bacterial genomes that contained a trypsin-MBL gene pair and for which a 16S rRNA sequence was available. Individual maximum-likelihood trees were constructed for trypsin domains, MBLs, and 16S rRNA genes using the methods described above. Pairwise evolutionary distances between genomes were derived from each phylogenetic tree, and Spearman’s rank correlation analysis was employed to evaluate the congruence between corresponding distance matrices.

### Plasmid construction for phage assay

For AVAST-MBL system from *Vibrio cholerae* (*Vc*AVAST-MBL), AVAST and MBL genes were cloned into a pBAD33 plasmid and a pET-28a derivative containing a pBAD promoter, respectively. For *Kp*Hachiman-MBL system, HamAB or its mutants was cloned into a pBAD33 plasmid, and MBL or its mutants was cloned into a pET-28a derivative. The two plasmids were co-transformed into *E.coli* MG1655 for subsequent phage assays.

### Phage plaque assay

All tested phages except T4 and T7 were obtained from CreatiPhage. Phage plaque assay was performed as previously described^[10]^. *E.coli* MG1655 strains harboring the following expression constructs were grown to exponential phase in LB medium supplemented with 50 μg/mL kanamycin and 34 μg/mL chloramphenicol: the *Vc*AVAST-MBL system, the *Kp*HamAB-MBL system, their mutants, or an empty vector control. For the phage assay, 400 μL of each bacterial culture was mixed with 8 mL of melted LB agar (0.5% w/v) supplemented with 50 μg/mL kanamycin, 34 μg/mL chloramphenicol, and 0.05% (w/v) L-arabinose. The mixture was poured into bottom agar layer and allowed to dry for 1 h at room temperature. Ten-fold serial dilutions of each of the 12 tested phages were prepared in LB broth. A 2.5 μL droplet of each dilution was then spotted onto the top agar layer to screening for phages susceptible to the *Kp*HamAB-MBL or *Vc*Avast-MBL systems. After overnight incubation at 37°C, plates were photographed. ΦCP-EC-23022 was selected for the subsequent phage plaque assays to elucidate the working mechanism of HamAB-MBL. All plaque assays were performed in biological triplicate.

### Liquid infection assay

The exponential-phase *E.coli* MG1655 culture harboring either the defense system or an empty vector was induced with L-arabinose at a final concentration of 0.15% (w/v) for the *Kp*HamAB-MBL system or 0.2% (w/v) for the *Vc*Avast-MBL system, followed by incubation at room temperature for 1 h. After incubation, 100 μL of the mixture was transferred to a 96-well plate. Then, 100 μL of a 10-fold serial dilution of phage (in LB supplemented with appropriate antibiotics) or 100 μL of LB alone was added to the mixture to achieve the desired MOI. Plates were incubated at 37°C overnight with shaking in a microplate reader, and OD_600_ was recorded every 15 min. Three biological replicates, each consisting of two technical replicates, were performed.

### Growth Curve Analysis

To test the cell toxicity induced by the trypsin-MBL systems, Hachiman-MBL and Ago-MBL were selected to perform growth curve analyses. The genes encoding trypsin•HamAB or trypsin•Ago (and their mutants) were cloned into the pACYC-Duet vector. In parallel, the genes encoding the MBL components of the three systems (and their mutants) were cloned into a pET-28a vector with IPTG inducible promoter. For each trypsin-MBL system, two plasmids were co-transformed into *E.coli* Bl21 (DE3). A single colony was inoculated into LB liquid medium (supplemnted with 50 μg/mL kanamycin and 34 μg/mL chloroamphenicol) and cultivated at 37°C until an OD_600_ of ∼ 0.35. Then, the culture was transferred to a 96-well plate, and protein expression was inducted by adding IPTG to a final concentration of 200 µM. The plates were incubated in a microplate reader at 37°C with shaking, and the absorbance at 600 nm was measured at 10-minute or 15-minute intervals overnight. All the experiments were performed in biological triplicate.

### Recombinant protein expression and purification

The gene encoding MBL was cloned into the pET28a vector with N-terminal 6×His tag. Trypsin•HamA and HamB sequences were cloned into the pCOLA-Duet-1 vector with a Twin-Strep-SUMO tag fused to the N-terminal of trypsin•HamA. Proteins were expressed in *E.coli* strain BL21 (DE3). The cell cultures were grown in LB medium supplemented with 50 μg/mL kanamycin at 37°C to an OD_600_ of ∼0.6, then induced with 0.3 mM IPTG and shifted to 18°C for 16 h. The cell pellets were harvested and lysed by sonication in lysis buffer (500 mM NaCl, 50 mM HEPES pH 8.0, 5% (v/v) glycerol, 0.1% (v/v) Tween-20, and 10 mM DTT). MBL were purified by Ni-NTA chromatography. Ni-NTA resin was washed with lysis buffer and proteins were eluted with elution buffer (500 mM NaCl, 50 mM HEPES pH 8.0, 5% glycerol, 300 mM imidazole). Trypsin•HamAB was purified by STarm Streptactin Beads 4FF resin. Strep resin was washed with lysis buffer and proteins were eluted with elution buffer (500 mM NaCl, 50 mM HEPES pH 8.0, 5% (v/v) glycerol, 2.5 mM D-biotin, 0.1% (v/v) Tween-20, and 10 mM DTT). The Twin-strep-SUMO tag of HamAB was cleaved by incubation with Ulp1 protease at 4°C overnight. Following cleavage, the proteins were further purified by size exclusion chromatography (SEC) using a HiLoad 16/600 Superdex 200 prep grade column (Cytiva) equilibrated in equilibration Buffer (500 mM NaCl, 50 mM HEPES pH 8.0, 0.05% (v/v) Tween-20, and 1 mM DTT). The purity was assessed by SDS-PAGE and the fractions of interest were pooled. The proteins were concentrated with a centrifugal filter (Amicon Ultra), flash-frozen in liquid nitrogen, and stored at –80°C until use.

### pNP-PC Assay

The phosphorylcholine esterase activity of MBL, trypsin•HamAB and trypsin•HamAB-MBL was measured using p-nitrophenyl-phosphorylcholine (pNP-PC, Sigma) as the substrate. Assays were performed at 37°C in a reaction buffer containing 200mM NaCl, 20 mM HEPES pH 8.0, 1 mM Fe(NH_4_)_2_(SO4)_2_, 5 mM DTT and 0.1 mM CaCl_2_, in a total volume of 200 μL. Prior to the reaction with the substrate, trypsin•HamAB was pre-incubated with 3′ overhang DNA for 10 min. Enzyme activity was measured by monitoring the increase in absorbance at 405 nm, which corresponds to the release of p-nitrophenyl (pNP).

### Nitrocefin Hydrolase Activity Assay

The Nitrocefin hydrolase activity of MBL, trypsin•HamAB, and trypsin•HamAB-MBL was determined by incubating the purified proteins with 100 μM Nitrocefin in a reaction buffer containing 200 mM NaCl, 20 mM HEPES pH 8.0, and 2 mM MgCl_2_ at 37°C for 30 min. Prior to the reaction, trypsin•HamAB was pre-incubated with 3’ overhang DNA for 10 min. Hydrolysis was monitored by measuring the absorbance at 495 nm using a microplate reader.

### ATPase Assay

The ATPase activity of trypsin•HamAB was analyzed using Malachite Green Phosphate Detection Kit (Beyotine) to quantify phosphate release. A panel of nucleic acid substrates—including ssRNA, ssDNA, dsDNA, 3′ overhang, 5′ overhang, and plasmid DNA—were used for substrate screening. Reactions were performed by incubating 50 nM purified trypsin•HamAB with either 100 nM DNA, 100 nM RNA, or 40 μg plasmid DNA in ATPase buffer (100 mM NaCl, 2 mM MgCl_2_, 20 mM HEPES pH 8.0, 0.1% (v/v) Tween-20) at 25°C for 15 min. The reaction was initiated by adding 0.1 mM ATP and allowed to proceed for 30 min. The samples were diluted to a suitable concentration within the linear range of the Kit. Malachite Green Reagent A/B was added to stop the reaction and progress color development at 25°C for 10 min. Absorbance at 630 nm was measured by microplate reader (Tecan SPARK). The experiment was performed with three biological replicates.

### DNA Unwinding Assay

Substrates used in unwinding assay were prepared by mixing the 5′ FAM labelled fluorescent strand with a 1.2-fold excess of its complementary non-fluorescent strand in annealling buffer (50 mM NaCl, 20 mM HEPES pH 7.5, and 5 mM MgCl_2_). The mixture was heated to 95°C for 5 min and then slowly cooled to 25°C over 1 h. To analyze the DNA unwinding activity of trypsin•HamAB, 100 nM of the annealed substrates was incubated with 50 nM trypsin•HamAB in reaction buffer (200 mM NaCl, 20 mM HEPES pH 7.5, 2 mM MgCl_2_, and 2 mM ATP or ADP) in the presence of 3 μM unlabeled competitor DNA at 37°C for 30 min. Following incubation, the reactions were subjected to electrophoresis with 6% Tris-glycine gel. The fluorescent signals were visualized using a Tanon MINI Space 3000 gel image analysis system.

### *In vitro* DNA and RNA Cleavage Assay

Cleavage activity of trypsin•HamAB-MBL was assessed by incubating purified trypsin•HamAB and MBL, each at 200 nM, with 200 nM 5′ FAM-labeled RNA/DNA substrate in a reaction buffer (100 mM NaCl, 20 mM HEPES pH 8.0, 2 mM ATP, and 2 mM MgCl₂) at 37°C for 30 min. The reaction was terminated by adding 2× loading dye (8 M urea, 5×TBE, and 0.025% bromophenol blue). Samples were then resolved by electrophoresis on 12% Urea-PAGE gels and visualized using a Tanon MINI Space 3000 gel image analysis system.

The cleaved MBL pulled down from culture co-expressing HamAB and MBL was subjected to DNA cleavage following the same procedure as above.

### *In vitro* Plasmid Cleavage Assay

For plasmid cleavage, 200 nM trypsin•HamAB and 200 nM MBL proteins were incubated with 400 ng of pUC19 plasmid in a reaction buffer containing 100 mM NaCl, 20 mM HEPES pH 8.0, 2mM ATP, and 2 mM MgCl₂ at 37 °C for 30 min. The reaction was terminated by phenol extraction using an equal volume of phenol. The samples were loaded onto 1% agarose gel and visualized using Tanon MINI Space 3000 gel image analysis system.

The cleaved MBL pulled down from culture co-expressing HamAB and MBL was subjected to plasmid cleavage following the same procedure as above.

### *In vitro* Cleavage Assay of MBL with Protease HamAB

The cleavage of MBL by protease HamAB and its mutants were assayed in a cleavage buffer containing 200 mM NaCl, 20 mM HEPES pH 8.0, 2 mM MgCl₂, 1 mM DTT, 5% (v/v) glycerol, and 2 mM ATP. Briefly, 5 μM trypsin•HamAB was first pre-incubated with 5 μM 3′ overhang DNA at 37°C for 15 min. Subsequently, 5 μM MBL was added, and the reaction mixture was incubated for an additional 15 min at 37°C. Reactions were terminated by the addition of SDS loading buffer. The samples were then resolved by electrophoresis on a 15% SDS-PAGE gel, stained with InstantBlue Coomassie dye, and visualized using a Tanon MINI Space 3000 gel image analysis system. For ATP inhibition assay, 2 mM ATP of the cleavage buffer were replaced with 0 mM, 1 mM, 3 mM, or 5 mM ATP; 5 mM AMP-PNP; or 5 mM ADP. All other steps were performed identically to the standard assay described above.

### *In vivo* Cleavage Assay of MBL with Protease HamAB

Plasmids encoding Twin-strep-*Kp*MBL (in pET28a) or its mutants and HamAB (in pCOLA-Duet-1) or its mutants were co-transformed into *E.coli* BL21 (DE3) cells. Transformed cultures were seeded into L-Broth medium supplemented with 50 μg/mL kanamycin and 34 μg/mL chloramphenicol and grown at 37°C to an OD_600_ of ∼0.35. Protein expression was then induced with 0.3 mM IPTG, and the temperature was shifted to 18°C for 16 h. Cells were harvested by centrifugation and lysed by sonication in lysis buffer containing 500 mM NaCl, 50 mM HEPES pH 8.0, 5% (v/v) glycerol, 0.1% (v/v) Tween-20, and 10 mM DTT. The clarified lysate was subjected to affinity purification using a Strep-Tactin affinity column. Purified proteins were denatured in SDS loading buffer, resolved by 15% SDS-PAGE gel, and stained with InstantBlue Coomassie dye. Protein bands were visualized using a Tanon MINI Space 3000 gel image analysis system.

### N /C-terminal Sequencing by Mass Spectrometry

Purified dMBL fragments were separated on a 15% SDS-PAGE gel and stained with InstantBlue Coomassie dye. The three target bands were excised and destained using 50% acetonitrile in 50 mM ammonium bicarbonate. The gel pieces were dehydrated with 100% acetonitrile, air-dried, and subjected to reduction and alkylation. Reduction was performed with 10 mM dithiothreitol (DTT) at 56 °C for 1 h, followed by dehydration. Alkylation was carried out using 20 mM iodoacetamide (IAM) in the dark for 1 h. The IAM solution was then removed, and the gel pieces were washed with destaining solution and 100% acetonitrile before drying. For enzymatic digestion, the gel pieces were rehydrated and added with 0.025 μg/μL rLys-C protease, incubated at 37 °C for 16 h. Peptides were extracted by adding 200 μL of extraction buffer (5% trifluoroacetic acid, 50% acetonitrile, 45% water) followed by incubation at 37 °C for 1 h. After sonication and centrifugation, the supernatant was collected. The extraction was repeated, and the combined supernatants were concentrated to dryness using a vacuum centrifugal concentrator at 45 °C. The peptides were desalted, dried, and reconstituted in sample dissolution solvent (0.1% formic acid, 2% acetonitrile). The resulting in-gel digested peptides were identified by chromatography-tandem mass spectrometry (LC-MS/MS), performed by Bio-Tech Pack Technology (Beijing, China).

### Proteolytic analysis of fluorescent RK-peptide

The purified 500nM trypsin•HamAB was mixed with 500nM 3’overhang in buffer containing 20mM HEPES pH 8.0, 200mM NaCl, 2mM MgCl_2_ and 2mM ATP. Reactions were then initiated by adding fluorescent RK-peptide to a final concentration of 20μM. After that, fluorescence emission at 490nm of the reactions was read using a SPARK Multi-Mode Reader (Tecan) after excitation at 340nm. The plates were incubated in a microplate reader at 37 °C with shaking, and measurements were taken at one-minute intervals for 30 min.

### *In vitro* reconstitution of HamAB-DNA Assemblies

To obtain trypsin•HamAB-DNA complex, 7 µM purified trypsin•HamAB was incubated with 10 µM 3′ overhang DNA in buffer containing 20 mM HEPES pH 8.0, 250 mM NaCl, 2 mM MgCl_2_, and 2 mM ATP. The sample was then resolved with Superdex 200 10/300 column. The fractions interested were pooled and concentrated with a centrifugal filter (Amicon Ultra).

**Cryogenic Electron Microscopy Sample Preparation and Data Collection** An aliquot of 3.5 μL HamA/B DNA complex at a concentration of 0.8 mg/mL was applied to glow-discharged Quantifoil holey gold grids (R 1.2/1.3, Au 300 mesh) followed by 15 s wait time and 4.5 s blot time with the blot force set to 2. Then, the grids plunge-frozen in liquid ethane using a Virtrobot Mark IV (Thermo Fisher). The Vitrobot chamber was maintained at close to 100% humidity and 22°C. Cryo-EM images were manually collected on a FEI Titan Krios G3i (Thermo Fisher) operated at 300 kV and equipped with a K3 Summit direct electron detector and Falcon 4i detector. All cryo-EM movies were recorded in counting mode with EPU.

### EM Data Processing

Images were processed in cryoSPARC (v.4.2) unless indicated otherwise. Movie frames were aligned and summed using Motioncor2. Contrast transfer (CTF) parameters were estimated for individual particles on each micrograph using cryoSPARC. After patch CTF estimation, micrographs with a resolution estimation worse than 6 Å were discarded. Blob picker was used to pick particles, using a circular diameter of 60 to 250 Å. Particle picks were inspected and particles with NCC scores below 0.25 were discarded. Subsequent 2D classification, 3D classification and 3D refinement were carried out using cryoSPARC. All refinements followed the gold-standard procedure, in which two half data sets were refined independently. The overall resolutions were estimated based on the gold-standard criterion of Fourier shell correlation (FSC) = 0.143. Local resolutions were calculated in cryoSPARC and visualized using ChimeraX^[11]^.

For *Kp*trypsin•HamAB DNA complex [monomer, dimer, trimer], 7,388 movies were recorded, and 6,686 images were kept after manual inspection. After 2D classification, 294,668 particles out of 10,550,525 auto-picking particles were retained and subjected for 3D reconstruction and 3D classification. To improve the resolution, 55,170 particles in a class were retained. Further 3D refinement, followed by non-uniform refinement, local refinement and post-processing, resulted in a 3.23 Å average resolution 3D map [monomer]. In addition, 88,815 particles belonging to a class from the initial 3D classification were selected for further 3D refinement and post-processing, resulting in a 3.22 Å average resolution 3D map [dimer]. Similarly, a 3.52 Å average resolution 3D map of [trimer] was generated using 50,060 particles through 3D refinement, non-uniform refinement, local refinement and post-processing.

### Model Building

The structures of HamA and HamB (accessions in GenBank: WP_171888367.1 and WP_023301564.1) were individually predicted using AlphaFold3^[12]^, and the best ranking AlphaFold3 predicted structures out of five predictions were chosen as models. The model was rigid-body fitted into the density using UCSF ChimeraX and manual rebuilding in Coot^[13, 14]^. Nucleotides and amino acid residues were mutated to reflect the true sequence of the construct. Models were subsequently refined with phenix.real_space_refine^[15]^, using reference to the starting structures and restraints on protein secondary structure. The quality of the structural model was checked using the MolProbity program in Phenix^[16]^.

---

**Figure S1.**
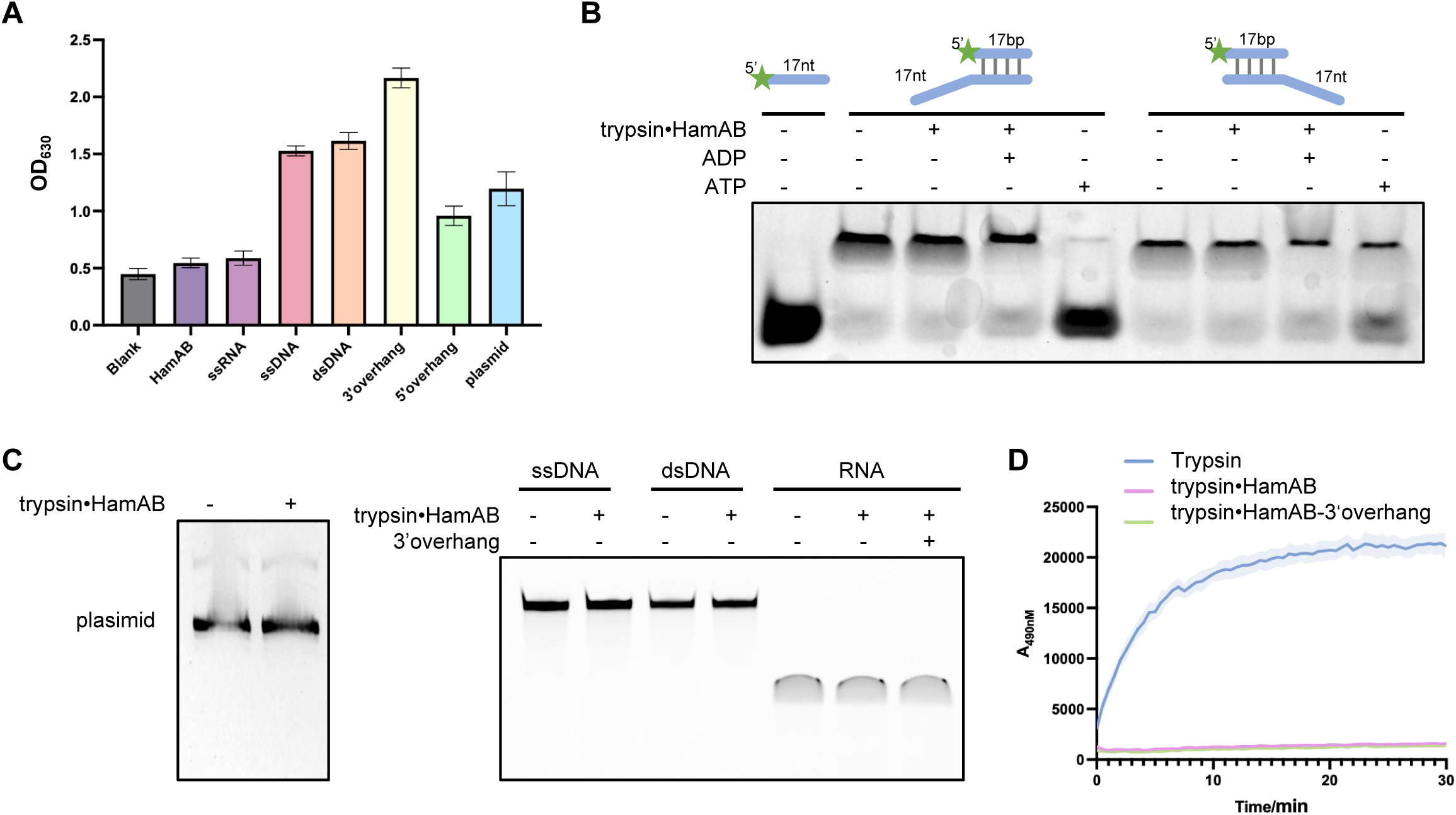
Functional characterization of trypsin•HamAB-MBL. **(A)** The ATPase activity analysis of trypsin•HamAB using Malachite Green. **(B)** Helicase activity analysis of trypsin•HamAB-MBL by DNA unwinding assay. **(C)** The nuclease activity analysis of trypsin•HamAB against plasmid (left gel), ssDNA, dsDNA, and RNA (right gel). **(D)** The protease activity of HamA using florescence-labeled peptide.

**Figure S2.**
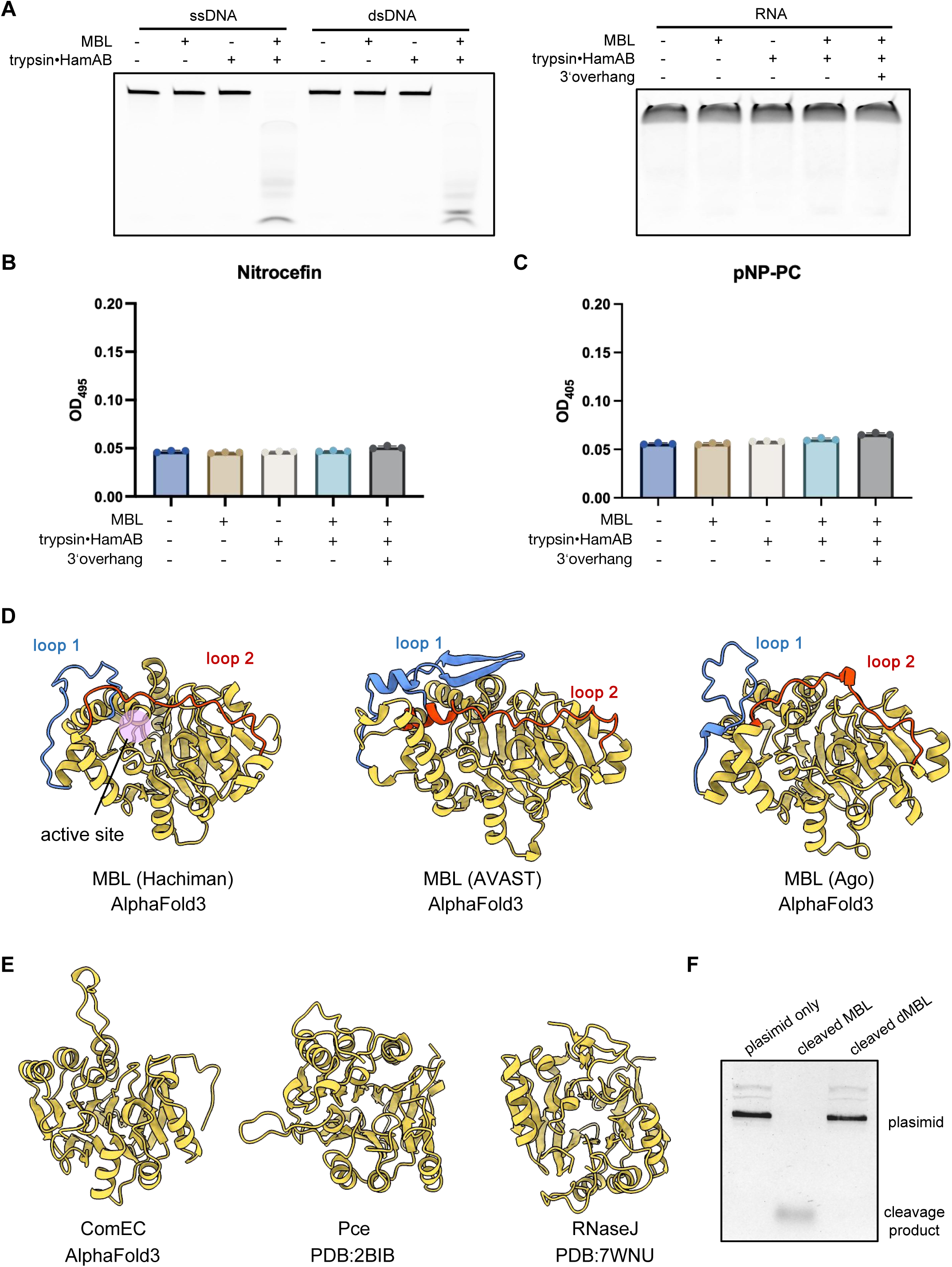
Function and structure characterization of MBL. **(A)** The nuclease activity analysis of trypsin•HamAB-MBL against ssDNA and dsDNA (left gel), and RNA (right gel). **(B)** The metal-β-lactamase activity analysis of MBL alone or trypsin•HamAB-MBL using nitrocefin as substrate. **(C)** The phosphorylcholine esterase activity analysis of MBL alone or trypsin•HamAB-MBL, using pNP-PC as substrate. **(D, E)** Structural comparison of MBLs (D) with the homologs (E) including ComEC, Pce, and RNaseJ reveals conservation of inhibitory loops in the defense systems containing trypsin-MBL module. The models of MBLs and ComEC are predicted by Alphafold3. **(F)** Plasmid cleavage assay of the cleaved MBL pulled down from culture co-expressing trypsin•HamAB and MBL.

**Figure S3.**
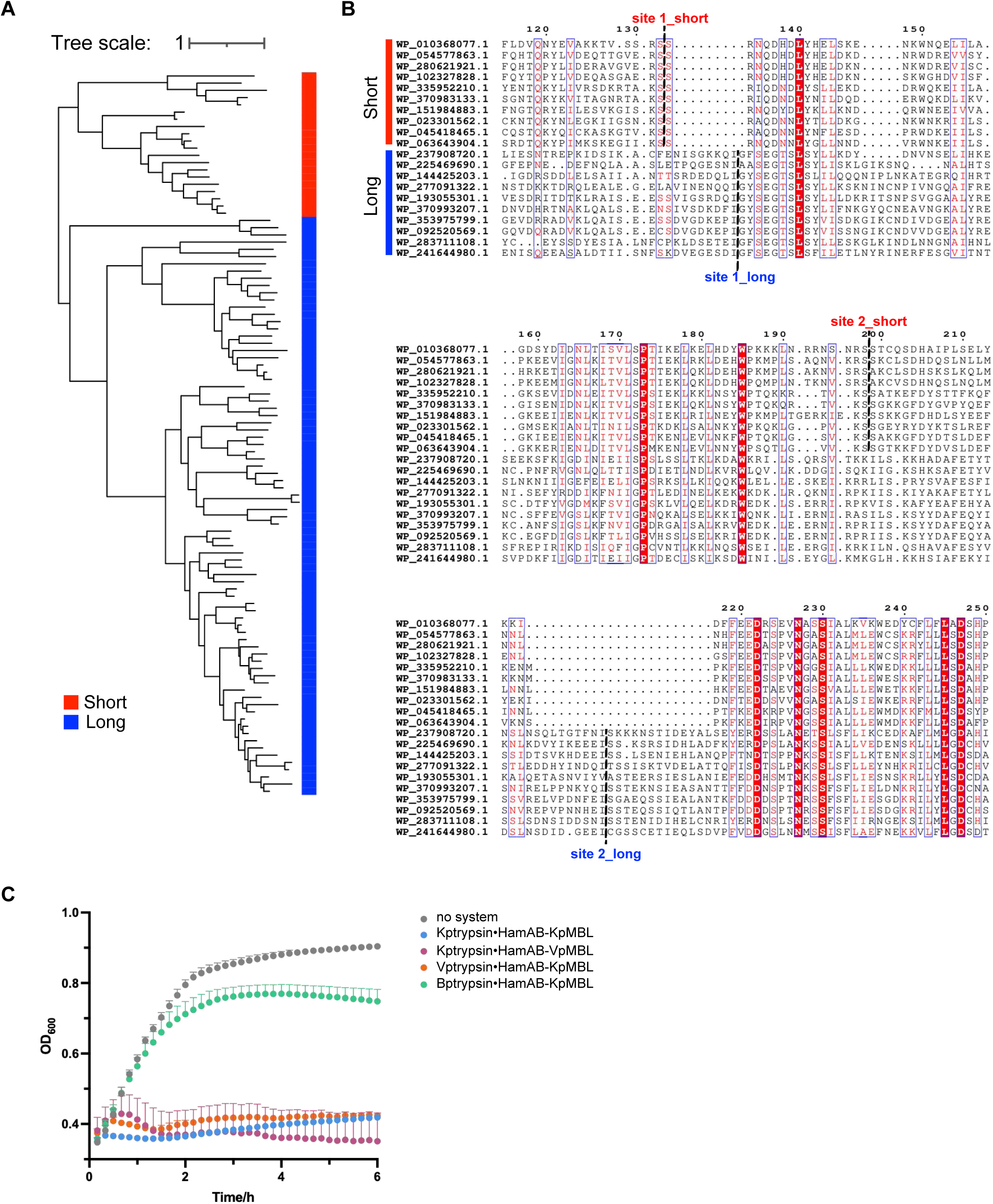
Classification of MBL from Hachiman-MBL systems according to the inhibitory loops. **(A)** Phylogenetic divergence of long- and short-form MBL proteins from Hachiman-MBL systems, shown in a subtree derived from Figure. 1A. **(B)** Sequence alignment of MBL from Hachiman-MBL systems. The two groups show divergence in length and cleavage sites of the inhibitory loops. The short and long groups are labeled with red and blue bar, respectively. **(C)** Growth curves of chimeric Hachiman-MBL systems exchanging long- and short-form MBLs.

**Figure S4.**
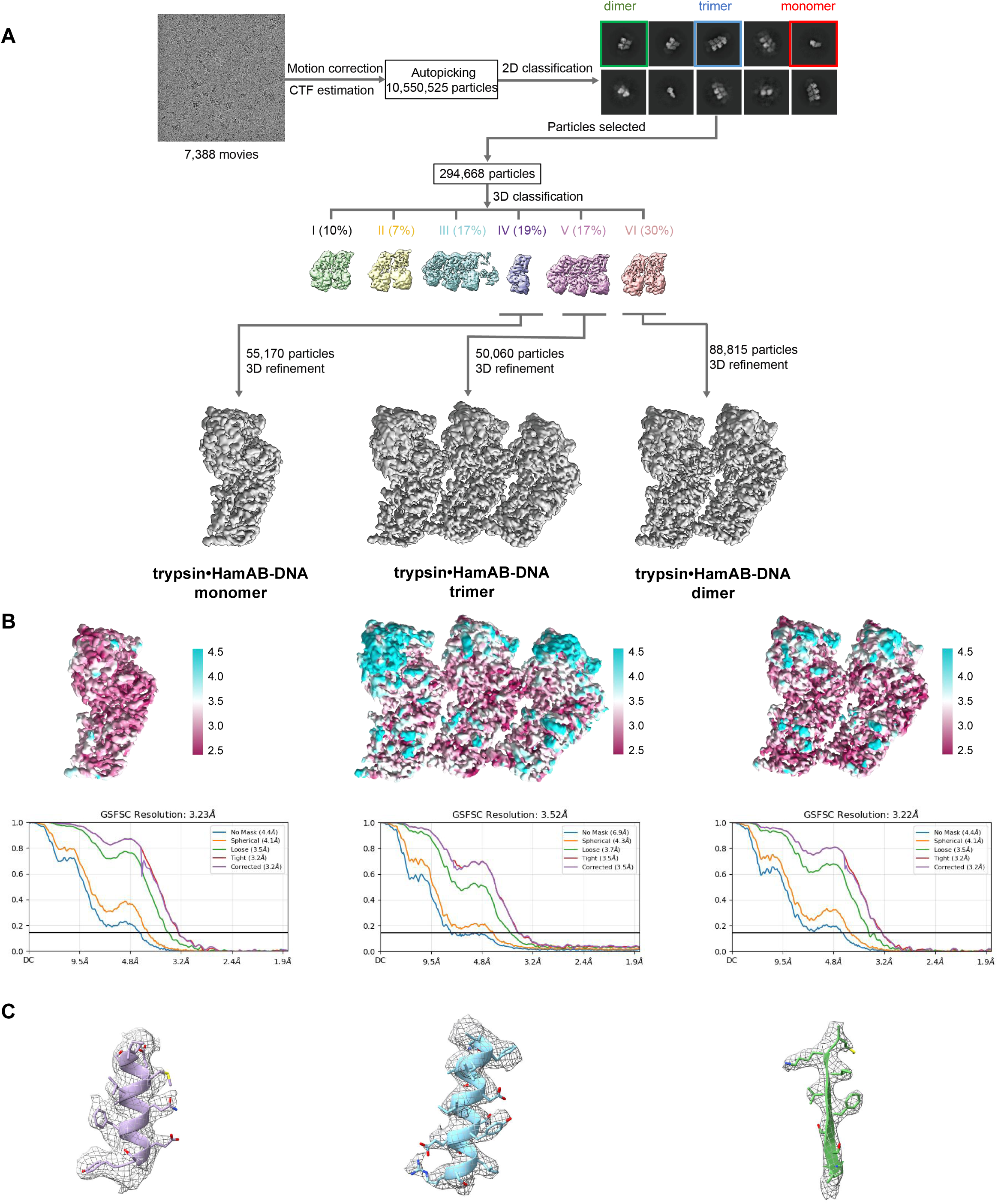
Single-particle cryo-EM analysis of trypsin•HamAB. **(A)** The flow-chat of single-particle cryo-EM analysis. **(B)** Cryo-EM density maps of trypsin•HamAB colored by local resolution, and gold-standard FSC of 0.143 resolution graphs is indicated for each complex. **(C)** The representative models and maps of trypsin•HamAB.

**Figure S5.**
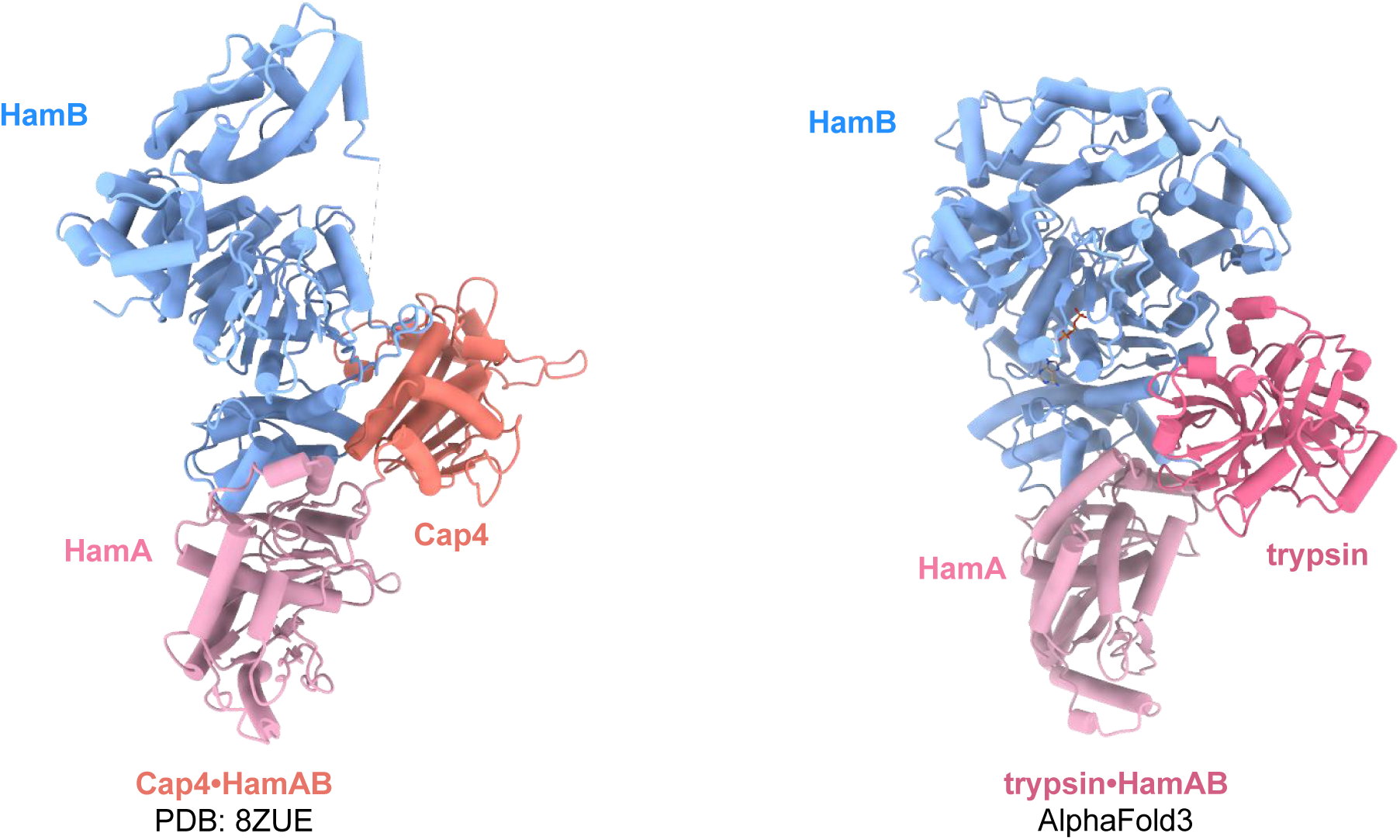
Structural comparison of trypsin•HamAB-ATP model with Cap4-HamAB structure.

