## Supplemental Table 2 for "The anti-phage mechanism of a widespread trypsin-MBL module"

### Supplementary information, Table S2 Cryo-EM data collection, refinement and validation statistics.

| | trypsin•HamAB-DNA, monomeric state (EMD-65964) (PDB 9WH1) | trypsin•HamAB-DNA, dimeric state (EMD-65968) (PDB 9WHK) | trypsin•HamAB-DNA, trimeric state (EMD-65977) (PDB 9WHU) |
|---|---|---|---|
| **Data collection and processing** | | | |
| Magnification | | 130 k | |
| Voltage (kV) | | 300 | |
| Electron exposure (e–/Å$^2$) | | 40 | |
| Defocus range (μm) | | -0.6 to -2.4 | |
| Pixel size (Å) | | 0.93 | |
| Symmetry imposed | | C1 | |
| Initial particle images (no.) | | 294,668 | |
| Final particle images (no.) | 55,170 | 88,815 | 50,060 |
| Map resolution (Å) | 3.2 | 3.2 | 3.5 |
| FSC threshold | 0.143 | 0.143 | 0.143 |
| Map resolution range (Å) | 2.0-39.0 | 2.0-40.0 | 2.0-44.0 |
| **Refinement** | | | |
| Initial model used (PDB code) | - | - | - |
| Model resolution (Å) | 3.4 | 3.6 | 3.9 |
| FSC threshold | 0.5 | 0.5 | 0.5 |
| Model composition | | | |
| Non-hydrogen atoms | 9486 | 18888 | 28353 |
| Protein residues | 1135 | 2270 | 3405 |
| Nucleotide residues | 10 | 16 | 25 |
| Ligands | 0 | 0 | 0 |
| *B* factors (mean, Å$^2$) | | | |
| Protein | 130.41 | 136.17 | 139.674 |
| Nucleotide | 188.55 | 193.47 | 189.46 |
| Ligand | - | - | - |
| R.m.s. deviations | | | |
| Bond lengths (Å) | 0.003 | 0.003 | 0.003 |
| Bond angles (°) | 0.587 | 0.601 | 0.643 |
| Validation | | | |
| MolProbity score | 2.50 | 2.66 | 2.66 |
| Clashscore | 10.02 | 13.37 | 14.30 |
| Poor rotamers (%) | 4.05 | 5.50 | 4.47 |
| Ramachandran plot | | | |
| Favored (%) | 90.98 | 92.66 | 91.48 |
| Allowed (%) | 8.66 | 7.21 | 8.05 |
| Disallowed (%) | 0.35 | 0.13 | 0.47 |
